# Proteome-wide screening for mitogen-activated protein kinase docking motifs and interactors

**DOI:** 10.1101/2021.08.20.457142

**Authors:** Guangda Shi, Jaylissa Torres Robles, Claire Song, Leonidas Salichos, Hua Jane Lou, Mark Gerstein, Benjamin E. Turk

## Abstract

Essential functions of mitogen-activated protein kinases (MAPKs) depend on their capacity to selectively phosphorylate a limited repertoire of substrates. MAPKs harbor a conserved groove located outside of the catalytic cleft that binds to short linear sequence motifs found in substrates and regulators. However, the weak and transient nature of these “docking” interactions poses a challenge to defining MAPK interactomes and associated sequence motifs. Here, we describe a yeast-based genetic screening pipeline to evaluate large collections of MAPK docking sequences in parallel. Using this platform we analyzed a combinatorial library based on the docking sequences from the MAPK kinases MKK6 and MKK7, defining features critical for binding to the stress-activated MAPKs JNK1 and p38α. We subsequently screened a library consisting of ~12,000 sequences from the human proteome, revealing a large number of MAPK-selective interactors, including many not conforming to previously defined docking motifs. Analysis of p38α/JNK1 exchange mutants identified specific docking groove residues mediating selective binding. Finally, we verified that docking sequences identified in the screen could function in substrate recruitment *in vitro* and in cultured cells. Collectively, these studies establish an approach for characterization of MAPK docking sequences and provide a resource for future investigation of signaling downstream of p38 and JNK MAP kinases.

## INTRODUCTION

Protein kinases function through phosphorylation of a limited number of effector substrates (*1*). Kinases achieve specificity in part through complementarity between the catalytic cleft and residues surrounding the site of phosphorylation. However, many kinases recognize minimal phosphorylation site motifs that are insufficient to confer protein-level selectivity, and closely related kinases with identical phosphorylation site specificity can nonetheless target distinct sets of substrates. A key example is the mitogen-activated protein kinases (MAPK) family, a group of Ser-Thr kinases conserved widely in eukaryotes (*2*). Canonical animal MAPKs, including the extracellular signal-regulated kinases (ERKs), c-Jun N-terminal kinases (JNKs) and p38 MAPKs (hereafter, p38), are positioned at the bottom of three-tiered kinase cascades activated in response to diverse cellular stimuli. The different MAPK subfamilies phosphorylate largely unique substrate repertoires to elicit distinct cellular responses, yet each targets a minimal Ser/Thr-Pro consensus sequence. MAPK specificity is at least in part driven by docking interactions, in which regions of the kinase outside of the catalytic cleft recruit substrates through binding sites distal from their sites of phosphorylation.

A major hub for MAPK docking interactions is the D-recruitment site (DRS), a conserved region of the catalytic domain that binds to substrates, scaffold proteins, MAPK kinases (MKKs) and MAPK phosphatases (MKPs) (*3–5*). Several highly conserved binding partners bind to the DRS via domains with intrinsic tertiary structure (*6*). However, the DRS more generally recognizes short linear motifs (SLiMs) termed D-sites found in unstructured regions of MAPK interactors (*7–10*). D-sites bind to MAPKs with moderate affinity (~100 nM – 30 μM), promoting transient kinase-substrate interactions and dynamic remodeling of signaling networks in response to stimuli (*7*). DRS engagement can also impact the conformation and dynamics of the MAPK catalytic domain, which may be important in promoting activation by MKKs and inactivation by MKPs (*11–13*).

The MAPK DRS forms a groove consisting of three adjacent hydrophobic pockets (designated the Φ_L_, Φ_A_ and Φ_B_ pockets) and a proximal negatively charged “common docking” (CD) site (*7, 8, 14, 15*). D-site sequences include a cluster of basic residues complementary to the CD site, a variable linker, and two or three hydrophobic (ϕ) residues (in a ϕ_L_-x-x-ϕ_A_-x-ϕ_B_, ϕ_L_-x-ϕ_A_-x-ϕ_B_, or ϕ_A_-x-ϕ_B_ arrangement) that engage the corresponding Φ pockets (*10, 15*). Additional sequence features within the context of this general motif appear to confer selectivity for particular MAPK subfamilies (*7, 9, 16, 17*). Structural studies of MAPK•D-site complexes have revealed distinct binding modes associated with these subfamily-selective motifs, driven by arrangement of the hydrophobic residues as well as the sequence composition and bound conformation of the D-site linker sequence (*7, 8, 11, 18–20*).

Knowledge of these MAPK-selective D-site motifs has driven computational approaches to identify novel MAPK substrates by scanning protein databases for sequences similar to previously established interactors (*9, 21*). However, experimental approaches for the discovery of additional MAPK-selective sequence motifs are needed to better define MAPK interactomes and to understand how they assemble. Here, we describe a genetically encoded library screening platform to identify new MAPK-interacting D-sites that exploits the conservation of the core cascade from humans to budding yeast. We used this strategy to analyze a saturation mutagenesis library of MKK-derived D-sites interacting with the JNK1 and p38α MAPKs and discovered previously unknown features of their corresponding motifs. We subsequently screened these MAPKs against a large library of human proteomic sequences, which allowed us to further refine the motifs and to identify new direct interaction partners. The screen differentiated between known p38α and JNK1 interactors and identified an extensive number of binding partners that are selective for either kinase. The dataset therefore provides an unbiased resource to probe selective MAPK signaling pathways and will be an important resource for future investigation into MAPK biology.

## RESULTS

### A yeast-based system for evaluation of MAPK docking sequences

We exploited the evolutionary conservation of core MAPK cascades to enable screening complex D-site sequence libraries in budding yeast. Yeast require the p38/JNK ortholog Hog1 for adaptation to osmotic stress (*22*). Normal activation of the Hog1 cascade by hyperosmolarity induces transient cell cycle arrest, but constitutive signaling causes stable growth suppression (*23*) (Fig. 1A). Presumably because it is deregulated in yeast, we found that co-expression of mammalian MKK6 with p38α was toxic to a strain lacking Hog1 and its cognate MKK Pbs2 (Fig. S1). MKK6 harbors a D-site located upstream of the catalytic domain (Fig. 1B) that promotes its phosphorylation of p38α (*24*). Because growth suppression in the context of WT p38α, which partially autophosphorylates in yeast, occurred independently of the MKK6 D-site (Fig. S1), we examined a fully MKK-dependent p38α mutant (L195A) (*25*). Growth impairment by p38α^L195A^ was less severe than for WT p38α, dependent on an intact MKK6 D-site, and partly alleviated by exchanging the MKK6 D-site with that of MKK7 (MKK6^D7^) (Fig. 1C). Furthermore, MKK6^D7^, but not WT MKK6 or MKK6^ΔD^, partly suppressed cell growth when co-expressed with the analogous JNK1^L198A^ mutant (Fig. 1D). These results provide a system in which yeast growth is controlled by a D-site interaction with either the p38α or JNK1 MAPK.

**Figure 1.**
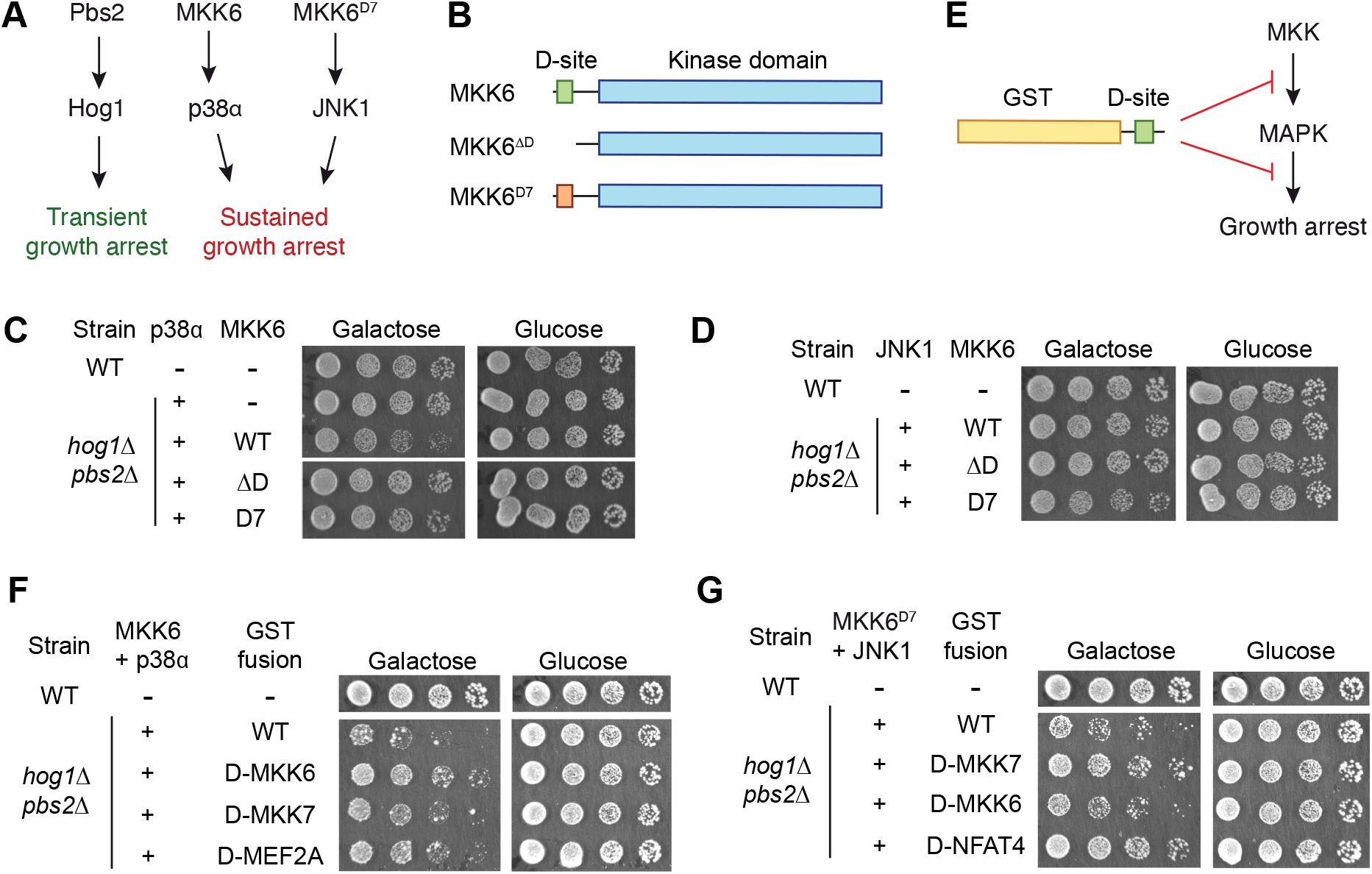
A system coupling MAPK D-site interactions to yeast cell growth. (A) Domain structure of MKK6 and D-site variants. MKK6^D7^ replaces the native p38-selective D-site with that from MKK7, which binds only to JNK. (A) Scheme showing the impact of replacing yeast MAPK pathway components with their human homologs on cell growth. (C) Assay showing growth inhibition of a *hog1Δ pbs2Δ* strain upon co-expression of p38α and the indicated MKK6 variants. Cells were grown in liquid culture, derepressed in raffinose media, and then spotted in 5-fold serial dilutions on solid media containing either galactose (to induce p38 expression) or glucose. (D) Growth assay of yeast co-expressing JNK1 and MKK6 variants conducted as in (C). (E) Scheme showing potential mechanisms for the growth rescue provided by expressing GST-D-site fusion proteins. (F) Effect of expressing cognate (MKK6, MEF2A) or non-cognate (MKK7) D-sites fused to GST on growth arrest mediated by p38α-MKK6 co-expression. Cells were grown and plated as in (C). (G) As (F) except with JNK1-MKK6^D7^ co-expression along with cognate (MKK7, NFAT4) or non-cognate (MKK6) D-sites.

In order to evaluate D-site sequences for their MAPK binding capacity, we introduced a third component into this system by ectopically expressing a D-site peptide. We reasoned that engagement of the MAPK DRS by the D-site would rescue growth inhibition by blocking either MAPK-MKK or MAPK-substrate binding (Fig. 1E). We first engineered *hog1Δ pbs2Δ* strains harboring chromosomally integrated cassettes for constitutive expression of the MKK (WT MKK6 or MKK6^D7^) and for inducible expression of the MAPK (p38α^L195A^ or JNK1^L198A^). We next introduced plasmids inducibly expressing D-site peptides fused to GST into these strains and assessed their growth under inducing or non-inducing conditions. Expression of p38α-binding D-site peptides derived from MKK6 or MEF2A both substantially reversed growth impairment associated with co-expression of MKK6 and p38α^L195A^, while the MKK7 D-peptide improved growth to a lesser extent (Fig. 1F). Likewise, D-site peptides from MKK7 and the JNK substrate NFAT4, but not from MKK6, rescued growth of the strain expressing MKK6^D7^ and JNK1^L198A^ (Fig. 1G). These experiments establish a system in which expression of a MAPK-binding D-site is coupled to cell growth, providing the basis for screens to identify sequences engaging the MAPK DRS.

### Comprehensive mutagenesis of MAPK docking sites

The importance of specific D-site residues in mediating selective interaction with MAPKs has been established by analysis of mutant proteins and synthetic peptides (*16, 17, 26, 27*), but low-throughput approaches can only analyze a limited number of variants. To more comprehensively analyze docking specificity using our yeast system, we designed a library of 715 unique sequences that included all possible single amino acid substitutions, as well as double Ala substitutions in all pairwise combinations, to D-sites from MKK6 and MKK7 (Fig. 2A, Data file S1). Oligonucleotides encoding these sequences were custom synthesized and cloned as a pool into the yeast GST fusion vector (Fig. 2B). Plasmid pools were introduced into MAPK/MKK-expressing yeast strains, and liquid cultures were expanded in derepressing (raffinose) media. A portion of the culture was reserved for sequencing, and the remainder was split and propagated under either inducing (raffinose + galactose) or non-inducing (glucose) conditions. At various times, cultures were sampled, plasmids extracted from cells, and the D-site region PCR amplified with barcoding primers. PCR samples were pooled and analyzed by Illumina sequencing, providing the relative abundance of each component of the library at each time point. While no sequences changed substantially (>10%) in relative abundance under non-inducing conditions, library representation became skewed upon induction of either MAPK (Fig. 2C,D, Fig. S2, Data file S2). The change of representation of each variant over time was fit to an exponential function to provide a measure of its relative depletion or enrichment within the screen (Fig. 2C,D, Data file S3).

**Figure 2.**
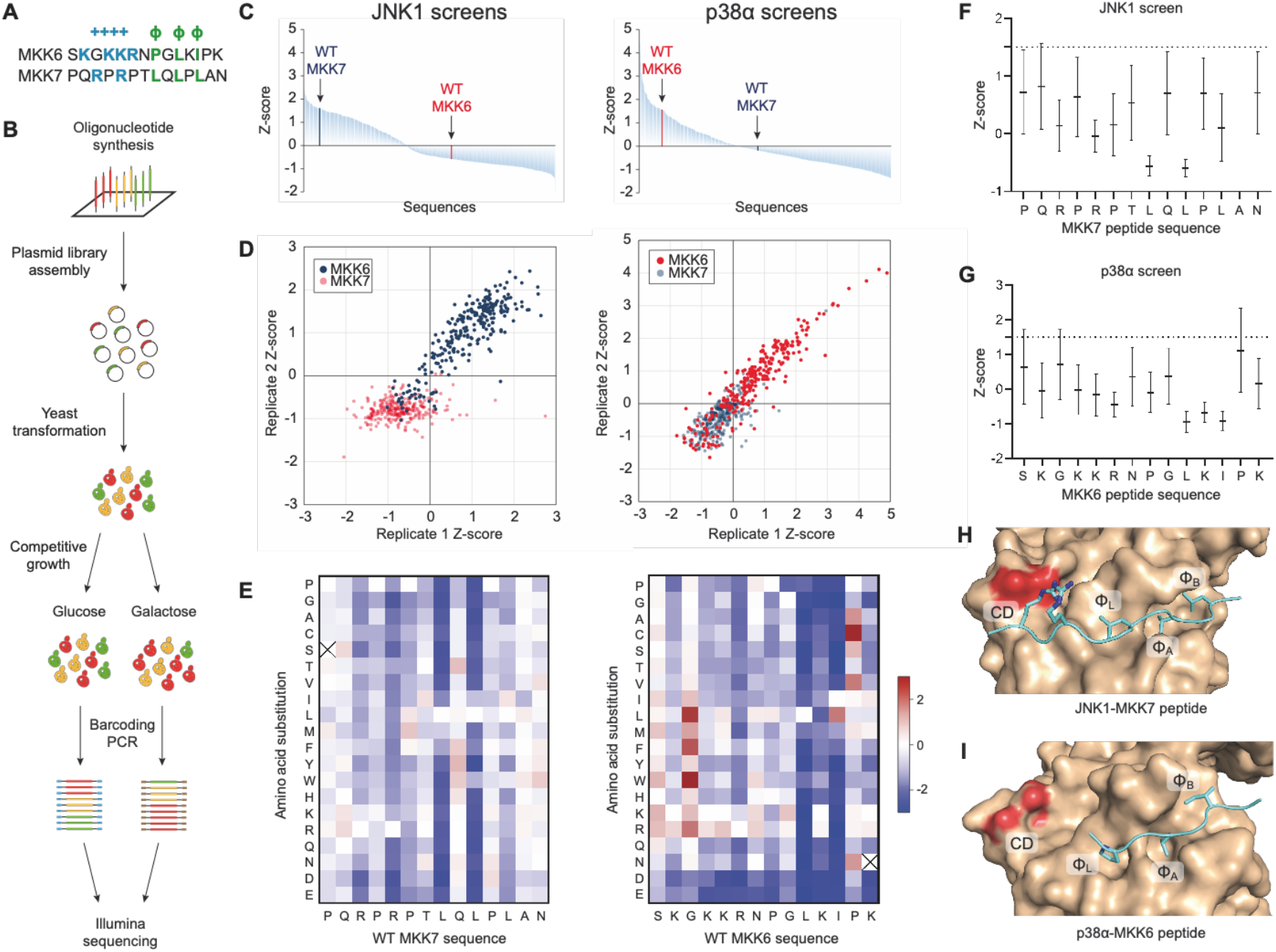
Combinatorial library screens. (A) A schematic showing the workflow of the screen. (B) Waterfall plots showing the average Z-score for the enrichment/depletion rate of each D-site sequence variant from two independent screens against JNK1 (left) or p38α (right). WT MKK6 and MKK7 D-sites are highlighted. Read counts from Illumina sequencing runs are provided in Data file S1. (C) Scatter plot showing correlation of Z-scores between two replicate screens for JNK (left) and p38α (right). Blue, MKK6 variants; red, MKK7 variants. (D) Heatmaps showing the effect of each amino acid substitution to the MKK6 D-site in the p38α screen (left) and MKK7 D-site in the JNK screen (right). Values are normalized to the respective WT sequence (white), with red indicating enrichment and blue indicating depletion of the sequence from the population. Crossmarked boxes indicate sequences missing from the screening library. (E) Graph showing mean normalized enrichment scores of all double alanine substitutions to each position in the MKK7 D-site sequence in the JNK1 screen. The dotted line indicates value of the WT sequence. Error bars show SD for the 12 variants at that position. (F) Data for MKK6 double Ala variants in the p38α screen are plotted as in (E). (G) Crystal structures of JNK1 (tan) in complex with the MKK7 D-site (cyan) with the CD region and three hydrophobic pockets indicated (PDB entry 4UX9) (*13*). (H) As for panel (G), but showing p38α bound to the MKK6 D-site (PDB entry 5ETF) (*28*).

As anticipated, variants derived from cognate D-sites (MKK6 for p38α; MKK7 for JNK1) were generally enriched during selection in two separate experiments, while with few exceptions non-cognate D-sites were depleted (Fig 2D). Most substitutions to cognate D-sites resulted in slower growth rates, suggesting that the WT sequences are largely optimal, but for both p38α and JNK1 some variants consistently conferred faster growth (Fig. 2E). For JNK1, Leu8 and Leu10, which engage the Φ_L_ and Φ_A_ pockets (*13*), appeared most critical for binding, while Leu12 that binds the Φ_B_ pocket was more tolerant to substitution (Fig 2E,F,H). As anticipated, substitutions to the N-terminally positioned basic residues, Arg3 and Arg5, that bind to the CD region, were also deleterious. For JNK1 the largest improvement was observed by aromatic substitutions to Gln9. A Tyr residue located at the same position in the NFAT4 D-site appears to provide intra- and intermolecular van der Waals contacts in the co-crystal structure with JNK1 (Fig. S3A) (*7*). For p38α, the most deleterious single (Fig. 2D) and double (Fig. 2F) amino acid substitutions were to Leu10 and Ile12, which engage the Φ_A_ and Φ_B_ pockets in the hydrophobic groove (Fig. 2I) (*7, 28*), where only hydrophobic residues were tolerated. As anticipated, substitutions to basic residues in the N-terminal cluster were also disfavored, and incorporation of additional Lys or Arg residues in this region appeared to promote binding. Surprisingly, mutating either Lys11 or Lys14 near the C-terminus, not previously reported as essential for interaction with p38α, also led to slower growth. Compared to JNK1, p38a was generally less tolerant of acidic residues, particularly in the C-terminal region (positions 11 - 14), likely related to a higher net negative charge in the DRS (see below). These observations suggest that p38α D-site binding may be in part driven by bulk electrostatics, rather than position-specific ionic interactions. At two sites near the N-terminus, binding was apparently improved by substitution with hydrophobic residues that may access an additional hydrophobic pocket (termed the Φ_U_ pocket) previously observed in some p38 and ERK D-site complexes (Fig. S3B) (*11, 28*). Only three residues within the MKK6 D-site sequence appeared suboptimal for binding, among them a Pro residue immediately downstream of the L-x-I motif. Overall, these positional scanning screens both confirmed elements of the MKK6 and MKK7 D-sites known to be critical for MAPK binding, and also identified additional features that favor or disfavor DRS interactions.

### Screening the human proteome for MAPK-interacting D-sites

The screens described above allowed us to define key features of the MKK6 and MKK7 D-sites. However, it is established that different D-site peptides can assume distinct binding modes at the DRS, each associated with its own conformation and associated sequence motif (*7, 9, 11, 13, 19*). To enable discovery of multiple MAPK-targeting motifs and to discover new p38 or JNK interactors, we designed a large library consisting of candidate D-site sequences derived from the human proteome following similar criteria to those used previously for *in silico* screens (*9, 21*) (Fig. 3A). We identified ~50,000 occurrences of the general D-site motif (defined as [RK]-x_0-2_-[RK]-x_3-5_-[ILV]-x-[FILMV]) in the human proteome. To increase the likelihood that sites would be accessible to bind MAPKs, we excluded sequences that fell within defined Pfam domains or were annotated to be extracellular or located within the ER or the Golgi apparatus. The remaining sequences were incorporated into a final library of 11,756 sequences from 5426 proteins, including most established JNK- and p38-interacting D-sites (Data file S1, Table S1).

**Figure 3.**
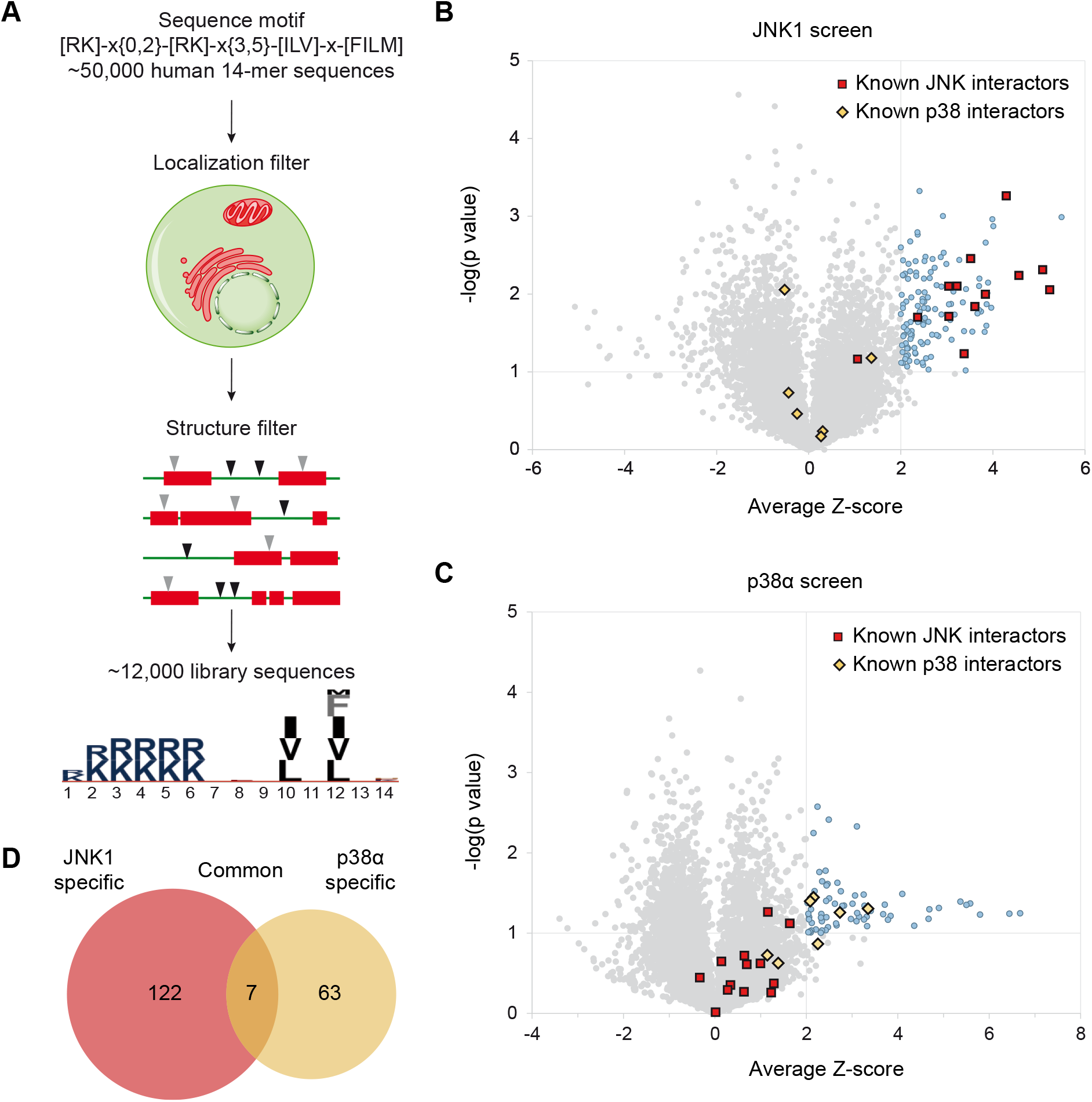
Proteomic library screens. (A) Schematic depicting selection of candidate D-site sequences from the human proteome. (B) and (C) Volcano plots for the JNK1 and p38α screens, respectively, showing mean Z-scores (n=3). The *p*-values (two-tailed heteroscedastic t-test) compare Z-scores for a given D-site with those of the entire population. Established functional JNK- and p38-interacting D-sites are indicated respectively as red squares and yellow diamonds. (D) Venn diagram showing overlap of hits (Z ≥ 2, *p* ≤ 0.1) from the two screens.

Coding sequences for all components of the library were introduced as a pool into the yeast GST fusion vector and screened for interaction with p38α and JNK1 as described above for the positional scanning libraries (Data file S4). Sequences were ranked by the average Z-score from three replicate screens (Data file S5). We observed established D-sites to be strongly enriched in screens for their respective MAPK (Fig 3B, data file S4). For JNK1, four of the five most highly ranked sequences (corresponding to JIP1, NFAT4, MKK4 and BMPR2) were previously reported docking sites. We defined “hits” as those sequences with an average Z score ≥ 2 and −log_10_(*p*-value) ≥ 1. By these criteria, all save one of the known JNK-interacting D-sites present in our library scored among the 133 hits in the JNK1 screen (Fig. 3B, Table S2). For p38α, we identified 71 hit sequences including four of seven known interacting D-sites (MKK4, MKK6, MEF2A and MEF2C). We note that the remaining D-sites (MKK3, PTPN5 and PTPN7) were also enriched in all screens but did not meet our hit threshold. We observed little overlap among hits for p38α and JNK1 with only seven sequences on both lists (Fig. 3C), and no non-cognate D-sites scored as hits for either MAPK. The strong enrichment of previously known D-sites suggests that other hits are likely to include authentic MAPK-interacting sequences.

In order to validate results from our screens, we examined the capacity of individual enriched sequences to selectively bind p38α and JNK1. To assess MAPK binding, we examined that ability of synthetic peptides to inhibit JNK1 and p38α activity toward a fluorogenic substrate incorporating a requisite D-site (*29*). We chose 24 hit sequences that had not previously been established to bind the corresponding MAPKs. As a measure of relative binding affinity, we determined IC_50_ values for inhibition of both kinases from assays conducted at a range of competitor peptide concentrations (Fig. 4A, Table S3). We found that all sequences scoring as hits in the JNK1 screen indeed bound JNK1 with higher affinity than p38α (Fig. 4A,B). While p38α was favored by most of its hit sequences, one of them was non-selective, and two slightly favored JNK1. We noted a tendency for JNK1 to bind D-sites more tightly than p38α, with five having IC_50_ values below 2 μM in comparison to only one for p38α (Fig 4A). This phenomenon may reflect an intrinsic capacity for JNK1 to bind tightly to docking sites or a more stringent affinity cutoff for enrichment in the yeast-based screen. Overall, these experiments confirm that most sequences selected in our screen bind selectively to p38α and JNK1 and do so with affinities comparable to established D-sites.

**Figure 4.**
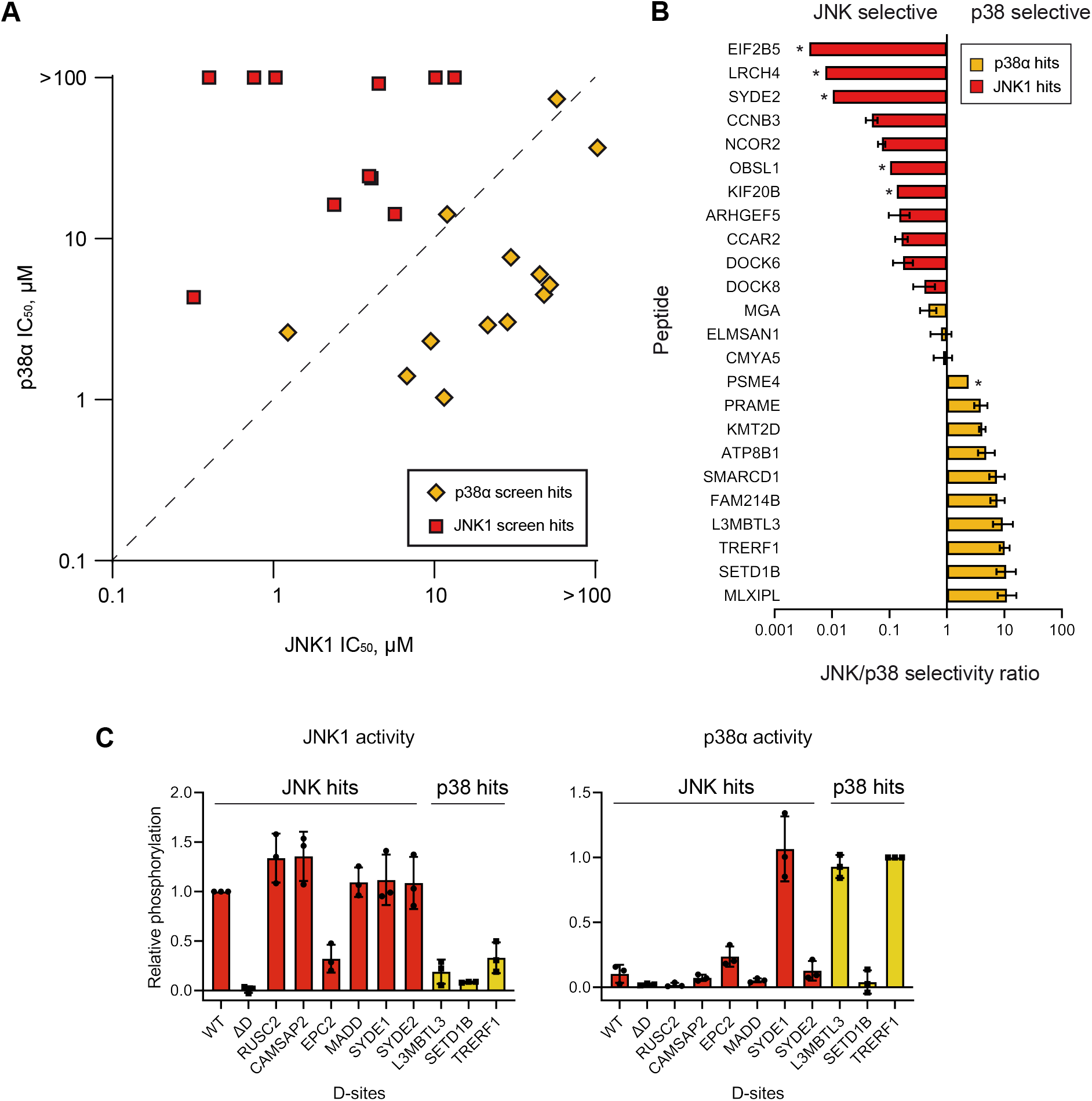
Hit validation. (A) Plot showing mean IC_50_ values (n = 3) for inhibition of the indicated MAPK by each of 25 chemically synthesized hit peptides. IC_50_ values too high to be confidently determined for p38α are indicated as being >100 μM and placed at the top edge. JNK1- and p38α-specific hits from the yeast screens are red squares and yellow diamonds, respectively. JNK1- and p38α-selective peptides fall respectively above and below the dotted line. (B) Graph shows ratios of JNK1 to p38α IC_50_ values for the indicated peptides, ordered from most JNK1-selective at top to most p38α-selective at bottom. For peptides binding weakly to p38α, the ratio was calculated using an IC50 value of 100 μM, and are indicated with an asterisk. Error bars show 95% confidence intervals. (C) Graphs show relative level of phosphorylation by JNK1 (left) and p38α (right) of chimeric NFAT4 constructs in which its D-site was replaced with those from the indicated hit proteins. Levels of phosphorylation were determined by immunoblotting with NFAT4 anti-phosphoSer165 antibody. Error bars indicate SD (n = 3).

As D-sites are often found in MAPK substrates, we further examined the capacity for selected p38 and JNK-targeting sequences to function in substrate recruitment. For these experiments we used an N-terminal fragment of the JNK substrate NFAT4 that harbors a D-site positioned upstream of established phosphorylation sites, including Ser165. We generated constructs in which the native NFAT4 D-site was either eliminated or substituted with hit sequences from the yeast screens. As anticipated, the WT fragment was robustly phosphorylated at Ser165 by JNK1 *in vitro*, and this was greatly reduced upon mutation of the docking site (Fig. 4C). Furthermore, substitution of four out of five JNK1 hit sequences tested fully restored Ser165 phosphorylation, while incorporation of p38α-targeting sites failed to do so. By contrast, p38α poorly phosphorylated the WT fragment, while incorporating either of two of its hit sequences (L3MBTL3 and TRERF1) converted NFAT4 to a p38α substrate. However, one other hit (SETD1B) failed to do so despite binding to p38α with comparable affinity (Table S3). This observation suggests that in some cases, D-site binding is insufficient to promote phosphorylation of an associated protein. D-site engagement can impact MAPK conformational dynamics to promote kinase activity (*30*), and our results suggest that such effects may occur in a sequence-specific manner. We also note that one D-site that was exclusively a JNK1 hit (SYDE1) promoted phosphorylation by both MAPKs and was presumably a false negative in the p38α screen.

### D-site sequence motifs conferring MAPK selective binding

Previous structural, biochemical, and computational analyses have identified multiple distinct sequence motifs selectively targeted by JNK and p38 MAPKs. Analysis of aligned hit sequences revealed that for both MAPKs, specific amino acids were significantly overrepresented at multiple positions in comparison to the full set of sequences in the library (Fig 5A). We expected that hit sequences for a given MAPK would conform to multiple distinct motifs that would not be evident from alignment of all hits. To deconvolute distinct motifs from the full dataset, we examined subsets of sequences in which a single overrepresented residue was fixed at one position (Fig. 5A). This analysis revealed two previously defined signatures within the JNK1 dataset: the “JIP class” motif (R-P-x-x-L-x-L) and the “NFAT4 class” motif (L-x-L-x-L/F) (*9*), with NFAT4 class sequences falling in one of two registers (with the first Leu residue occupying either position 8 or position 10 in the library). Among these core motifs, we observed specific residues to be enriched at intervening positions, for example hydrophilic residues downstream of the conserved Pro in the JIP class motif, and acidic/amidic residues intervening the ϕ_A_ and ϕ_B_ residues of both motifs. These two motif classes account for 82% of JNK1 hits, with the remaining sequences being enriched for a Leu residue in position 7 (producing a L-x-x-L-x-L motif) that likely occupies the Φ_L_ pocket similar to the Pro residue in the JIP class motif. Hit sequences for p38α were most strongly enriched for the previously defined “MEF2A class” motif (P/L-x-L-x-I/L-P) (*9*), with about 60% harboring a Pro or aliphatic residue in position 8 (ϕ_L_) (Fig. 5B). Of the remaining sequences, about half had a hydrophobic residue in position 7, and the other half were characterized by one or more basic residues upstream of the ϕ_A_ (position 10) Leu residue. These residual sequences also lacked selectivity for an Ile residue at the ϕ_B_ position or a Pro residue at position 13, suggestive of a distinct binding mode. In keeping with the results from the positional scanning library above, proteomic sequences selected by p38α were generally enriched for basic residues in positions proximal to the hydrophobic residues, a feature not generally considered as part of known interaction motifs. We note that there was no significant selection for Lys or Arg residues near the D-site N-terminus by either MAPK (Fig. 5A,B). However, because our library design included at least two basic residues in all sequences, they may still promote binding in a manner independently of their precise position.

**Figure 5.**
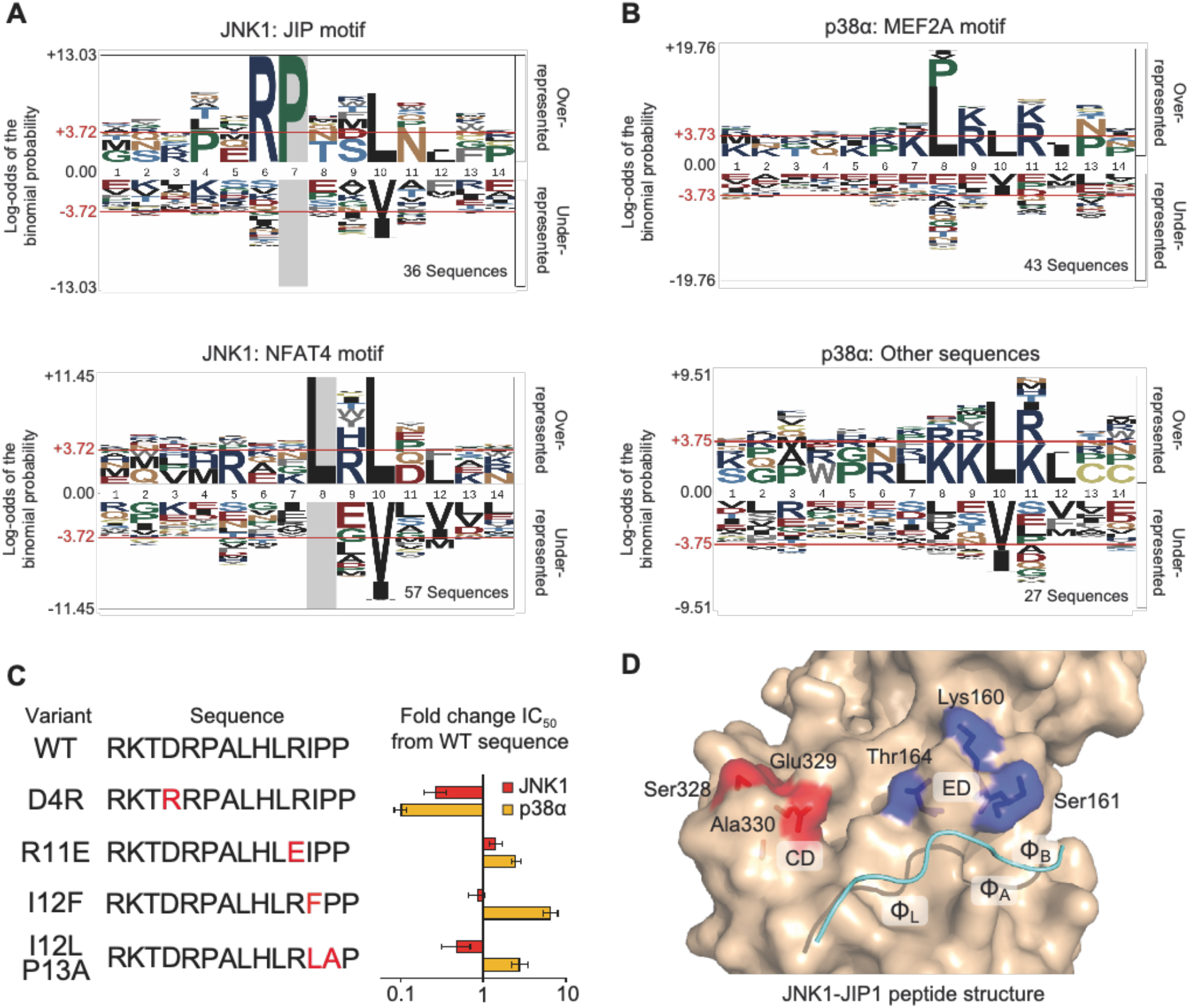
MAPK-interacting sequence motifs. (A) JNK1 hit sequences represented by pLogos(*62*). Top, subset of JNK1 hits having a Pro residue at position 7 (36 sequences) corresponding to the JIP class motif. Bottom, JNK1 hits with a Leu residue at position 8 (57 sequences) comprising the NFAT4 class motif. (B) Top, MEF2A motif class sequences including all p38a with either an Ile, Val, Leu, or Pro residue at position 8 or a Pro residue at position 13 (43 sequences). Bottom, all remaining p38a hits (27 sequences). (C) Peptide substitution analysis. Synthetic peptides corresponding to the p38α hit KMT2D peptide and the indicated variants were evaluated for competitive inhibition of JNK1 or p38α *in vitro*. Graph shows the average (n = 3) ratio of the IC_50_ value for the indicated point-substituted peptide to that of the WT peptide. Error bars show 95% confidence intervals. (D) Potential D-site specificity-determining residues. Two patches of surface residues in the docking groove of JNK, the CD (red) and ED (blue) regions, are shown mapped on the crystal structure of JNK1 in complex with the JIP1 peptide (PDB code 1UKI) (*18*).

We probed the importance of key elements of these motifs by substitution analysis of a p38α hit peptide derived from KMT2D (RKTDRPALHLRIPP) in assays of kinase inhibition as described above (Fig. 5C). Consistent with our screens and previous reports that the identity of hydrophobic residues can drive MAPK specificity (*16*), we found that substitution of the ϕ_B_ Ile residue with Phe led to a large decrease in p38α binding, while modestly improving binding to JNK1. Combined substitution of the Ile12-Pro13 sequence with Leu-Ala also made the peptide more JNK1 selective, suggesting a switch to an NFAT4 motif class. Substitution of Arg12 with Glu had a larger impact on binding to p38α in comparison to JNK, consistent with results from both the proteomic and positional scanning screens. Replacement of Asp4 with Arg improved binding to both MAPKs, confirming the importance of basic residues in that region. Collectively, these assays verify key sequence features selected by the two MAPKs in our screens.

Knowledge of a protein interaction motif can be used to discover new interacting proteins by searching databases for matching sequences (*31*). While our screening approach identifies MAPK interacting sequences present in the human proteome, authentic MAPK D-sites that do not conform strictly to the criteria used to build the library will be excluded. To identify D-sites not present within our screening library, we scanned the human proteome for sequences similar to our hits using the web application PSSMsearch (*32*). We performed separate searches using hit sequences corresponding to various p38α and JNK1 targeting motifs (Table S4). As anticipated, these searches returned sequences from the yeast screens that were used to build the PSSM, as well as sequences that were absent from the library. We chose 8 peptides excluded from the original library and evaluated their affinities for p38α and JNK1 in the competitive kinase assay (Table 1). With one exception, peptides bound to their predicted kinases with affinities comparable to those from the yeast screens (IC_50_ range 0.3 - 5.6 μM). These results confirm our ability to computationally identify new docking sequences, providing additional validation for our experimentally determined motifs.

**Table 1.**
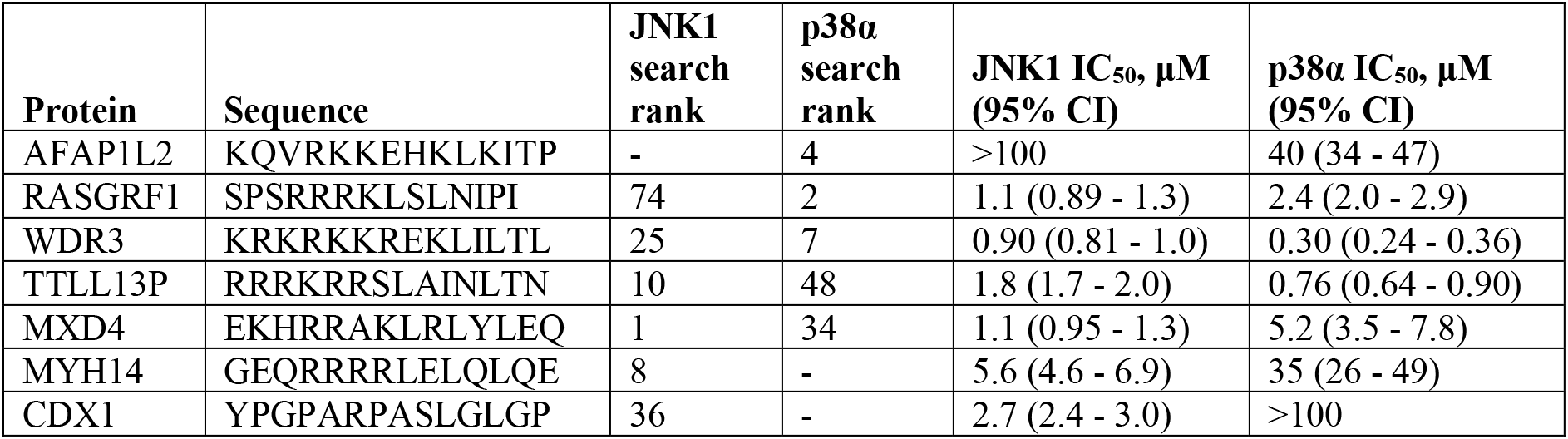
Inhibitory potency of peptide sequences identified by PSSMsearch.

### Determinants of MAPK-selective D-site interactions

We next considered features of the p38 and JNK DRS that might mediate selective targeting of distinct motifs. As noted above, p38α was more generally selective for basic residues throughout the D-site sequence. Notably, the p38α DRS is more negatively charged than that of JNK1, having nine acidic residues in comparison to four for JNK1. Unique acidic residues in p38α cluster at two sites: the so-called ED region proximal to the Φ_L_ and Φ_A_ pockets (*4*) and the CD region (Fig 5D). To examine the importance of these two regions to D-site specificity, we generated point mutants exchanging residues between p38α and JNK1 and examined binding of peptides corresponding to MAPK-selective motifs by competitive kinase assay (Table 2). Mutating either the ED or CD regions of JNK1 to the corresponding residues in p38α reduced binding of the JNK-selective NFAT4 and JIP1 peptides by an order of magnitude. While the JNK1 CD mutant had modestly increased affinity for the p38-selective MKK6 (MEF2A motif) and SETD1B (non-MEF2A motif) peptides, a larger effect was seen with the ED mutant, which had affinities similar to WT p38α for the two peptides. Conversely, the p38α ED mutant decreased the affinity of both cognate peptides to levels comparable to those seen for JNK1, while the CD mutant was without significant effect. None of the p38α mutants conferred detectable binding to the JNK1 cognate NFAT4 and JIP1 peptides. These experiments substantiate a role for a specific cluster of residues in mediating selective binding of D-sites to p38α, either through specific side chain interactions or though bulk electrostatic effects.

**Table 2.**
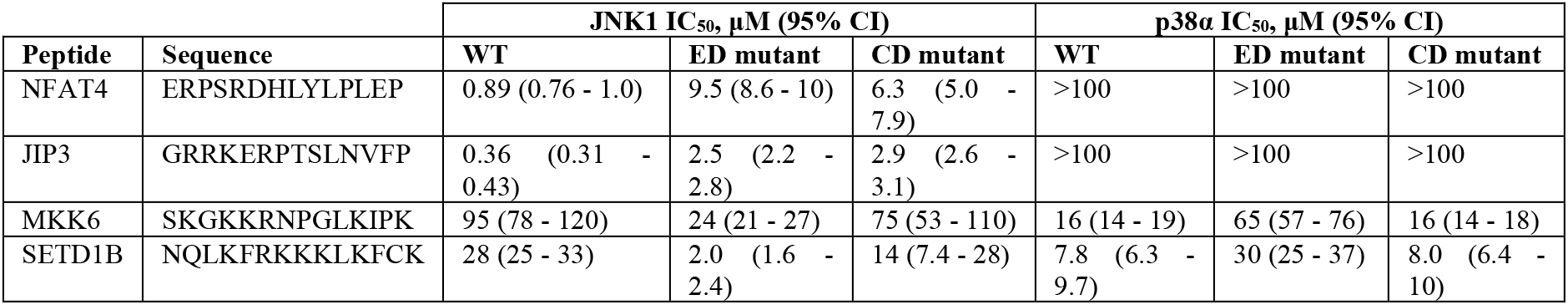
Impact of p38α/JNK1 exchange mutagenesis on peptide binding affinities. JNK1 ED mutant, K160N/S161E/T164E; JNK1 CD mutant, S328D/E329D/A330E; p38α ED mutant, N159K/E160S/E163T; p38α CD mutant, D315S/D316E/E317A.

### Discovery of docking-dependent MAPK substrates

To explore how our D-site screens might contribute to our understanding of MAPK function, we first performed gene set enrichment analysis (*33*) to identify potential cellular processes invovling p38 and JNK (Table 3). Hits for p38α were enriched for proteins involved in regulation of chromatin organization and gene transcription, with most (65%) localizing to the nucleus. These observations are consistent with nuclear translocalization of activated p38 and for its established roles in transcriptional regulation (*34*), and suggest broader control of gene transcription than that has been previously appreciated. While the observed enrichment of JNK1 hits for proteins involved in the JNK MAPK cascade was expected due to the abundance of known interactors, this category also included four upstream regulators not previously known to interact with JNK (TNIK, NCOR1, MAGI3, CARD9) that could constitute points for feedback regulation. In addition, JNK1 interactors were significantly enriched for cytoskeleton-associated proteins and regulators of signaling by small GTPases.

**Table 3.** Gene set enrichment analysis of p38α and JNK1 hits. FDR, false discovery rate.

| JNK1 screen | | p38α screen | |
| --- | --- | --- | --- |
| GO category | FDR q-value | GO category | FDR q-value |
| Regulation of Intracellular Signal Transduction | 1.44E-10 | Chromatin Organization | 9.80E-09 |
| Nucleoside Triphosphatase Regulator Activity | 5.82E-10 | Chromosome Organization | 2.73E-08 |
| Cytoskeleton Organization | 1.65E-09 | Catalytic Complex | 1.19E-07 |
| Small GTPase Mediated Signal Transduction | 2.85E-09 | Positive Regulation of Biosynthetic Process | 3.54E-07 |
| Guanyl Nucleotide Exchange Factor Activity | 7.90E-09 | Positive Regulation of Nucleobase Containing Compound Metabolic Process | 8.98E-07 |
| Positive Regulation of Catalytic Activity | 8.41E-09 | Nuclear Protein Containing Complex | 8.98E-07 |

Because D-sites can play a role in MAPK substrate recruitment, we reasoned that hits from our yeast screens include previously undescribed substrates. To verify that this was the case, we chose a pair of candidate substrates from our JNK1 screens, SYDE1 and SYDE2, that not previously known to interact with the kinase. SYDE1 and SYDE2 are GTPase activating proteins (GAPs) for RHO family small GTPases, with ascribed roles in embryonic development and neuronal function (*35, 36*). The JNK1-selected D-site sequences of SYDE1 and SYDE2 are found in an analogous position downstream of the GAP domain, though they belong to distinct motif classes (JIP and NFAT4-type, respectively). We found that JNK1 could phosphorylate purified SYDE1 and SYDE2 *in vitro* (Fig. 6A,B). Addition of a peptide corresponding to the NFAT4 D-site inhibited JNK1 activity toward both proteins, suggesting that phosphorylation was dependent on an interaction with the DRS. To determine whether they could be phosphorylated by JNK in cells, we transfected HEK293T cells with plasmids encoding either protein and treated cells with the protein synthesis inhibitor anisomycin to activate the JNK pathway in the presence or absence of the selective JNK inhibitor JNK-IN-8, and cell lysates were analyzed by Phos-tag SDS-PAGE and immunoblotting. We found that anisomycin induced phosphorylation of both proteins as judged by an electrophoretic mobility shift, which was reversed by treatment with the JNK inhibitor (Fig. 6C,D). Analysis of SYDE2 by liquid chromatography-tandem mass spectrometry (LC-MS/MS) revealed that a phosphorylation at a single residue (Ser1203) increased with anisomycin and decreased with JNK inhibitor treatment (Fig. 6E). This site is located 15 residues downstream of the SYDE2 D-site, consistent with prior observations that D-sites frequently direct MAPKs to phosphorylation sites proximal and downstream of the D-site (*37*). Overall, these observations verify that our ability to identify new MAPK substrates from our D-site screens.

**Figure 6.**
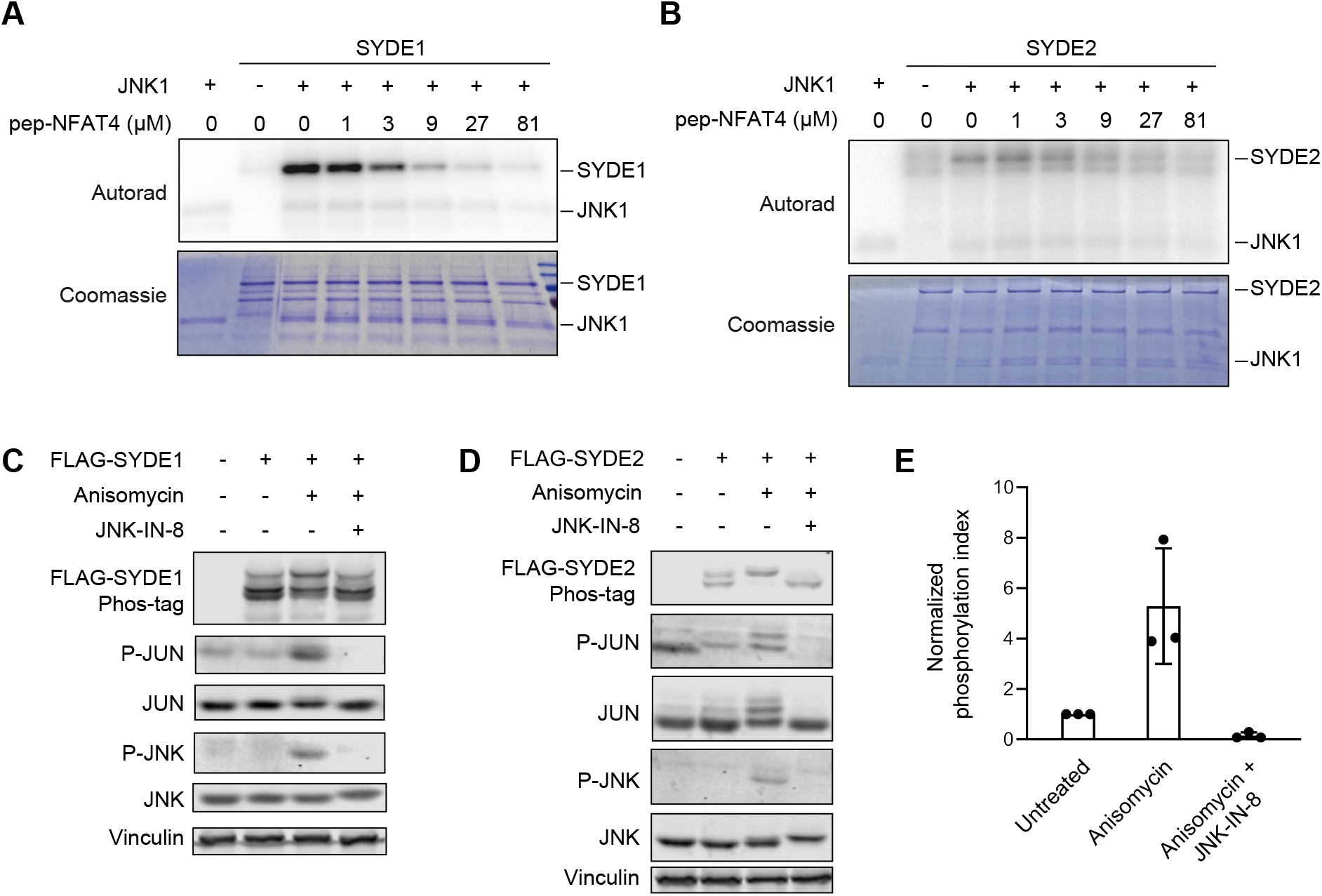
JNK substrate discovery. (A) *In vitro* radiolabel kinase assay showing JNK1 phosphorylation of full length SYDE1. Reactions were performed with increasing concentrations of the NFAT4 D-site competitor peptide to selectively block docking interactions. Representative images from three separate experiments are shown. (B) Kinase assay showing JNK1 phosphorylation of SYDE2, performed as in (A). (C) HEK293T cells transfected with a plasmid expressing FLAG-SYDE1 were treated with anisomycin following preincubation with or without the covalent JNK inhibitor JNK-IN-8. Lysates were subjected to either Phos-tag (top image) or standard (all others) SDS-PAGE, and immunoblotted with the indicated antibodies. (D) As in (C), except cells were transfected with a FLAG-SYDE2 expression vector. (E) MS analysis of SYDE2 phosphorylation. HEK293T cells transiently expressing FLAG-SYDE2 were treated as indicated. SYDE2 was immunoaffinity purified, and tryptic peptides were analyzed by LC-MS/MS. Graph shows the level of the Ser1082-phosphorylated peptide normalized to the total abundance of the corresponding peptide.

## DISCUSSION

Here we have described a genetic screening approach to the identification of kinase docking motifs and interacting sequences. Most prior screens for docking sequences have focused on protein phosphatases, which are generally described as lacking dephosphorylation site specificity and are hence dependent on non-catalytic interactions for substrate recruitment (*38–41*). However, there has been growing appreciation for the importance of non-catalytic SLiM-mediated interactions to kinase substrate targeting. For MAPKs, docking interactions can enforce selective targeting to individual subfamilies and can restrict the kinase to phosphorylate specific Ser or Thr residues in a given substrate(*9, 10, 37*). In other systems, docking is not absolutely required for phosphorylation *in vivo*, yet tuning the strength of such interactions can reportedly set phosphorylation rate. For example, SLiM-mediated recruitment to cyclin-dependent kinases through the cyclin subunit substrates controls the timing of phosphorylation within the cell division cycle (*42, 43*). In the case of the yeast LATS/NDR kinase Cbk1, an optimal docking sequence is not absolutely required for substrate phosphorylation yet confers robustness to perturbations that attenuate kinase activity (*44*).

Previous approaches to identify SLiMs mediating protein-protein or protein-enzyme interactions have used libraries of synthetic peptides or genetically-encoded phage display and cell surface display libraries (*40, 45, 46*). Our method involving reconstitution of signaling pathways in yeast involves tunable pathway inhibition through a competitive interaction between kinases in the MAPK cascade or with downstream effectors. This approach may be advantageous in that interactions occur within a eukaryotic cell while maintaining sufficiently high throughput to enable extraction of binding motifs. It also enables discovery of interaction partners that might escape detection in MS-based proteomics experiments due to low abundance or restricted patterns of expression. The expression level of D-site fusion proteins in yeast sets a relatively stringent affinity threshold providing a low false positive rate, but we may consequently fail to identify low affinity interactions. For example, while our approach selected almost all known JNK interactors in our library, it failed to identify the established D-site in the transcription factor ATF2. This is consistent with a recent report that regions of ATF2 outside of its SLiM make an additional contact with MAPKs to increase binding affinity (*47*). Likewise, the full p38α-interacting region of the phosphatase PTPN7/HePTP includes additional sequence flanking the canonical D-site (*48, 49*), and while enriched in our screen it fell below our hit threshold.

In this study we performed both a comprehensive mutagenesis screen of known docking sites as well as a screen of proteome-derived sequences. Screening positional scanning libraries has facilitated discovery of new kinase substrates conforming closely to the resulting motifs (*50*). Selection of proteomic libraries has an advantage in that it directly nominates candidate interaction partners and can discover high affinity sequences that might appear suboptimal. Furthermore, a sufficiently large set of interacting sequences will also provide key sequence features constituting the interaction motifs. Indeed, using our motifs we were able to identify additional high affinity JNK1 and p38α-binding sequences absent from our screening library. Our studies used a focused library built around a minimal consensus sequence shared by most known D-sites, excluding sites that bind in a reversed N-C orientation such as those found in MAPKAPKs (*7*). In the future we plan to adapt our approach to screen more complex “disorderome” libraries (*51*) that are not limited by a particular mode of interaction and may facilitate discovery of additional interaction motifs.

Several previous studies have used computational approaches to scan proteomes for MAPK-interacting D-sites based on consensus motifs defined by alignment of known interactors and by integrating structural constraints based on crystallographic studies of MAPK-D-site complexes (*9, 21*). While we did observe substantial overlap in our datasets, the majority of our hit sequences were not previously predicted (see Table S2). For example, 36% of JNK1 and 34% of our p38α hits were also discovered in the structure-guided *in silico* screens conducted by Remenyi and co-workers (*9*), and 11% of JNK1 hits had been identified by Bardwell and co-workers through database searches for sequences similar to known D-sites (*21*). We note that hits unique to our dataset were often dissimilar in sequence to those for which structural information is available. In particular over 90% of our p38α hits lacking an ϕ_L_ residue had not been previously predicted. These observations underscore the value of unbiased screens of proteome-derived libraries in the discovery of new interactors.

JNK and p38 MAPKs were originally identified as stress-activated kinases and are now known to have diverse roles in normal and disease physiology (*34, 52, 53*). We found significant enrichment for specific processes associated with proteins harboring MAPK-interacting D-sites uncovered in our studies, suggestive of expanded roles for JNK and p38. For example, p38 is a well-established regulator of gene transcription, directly phosphorylating more than a dozen sequence-specific transcription factors (*34*). In addition to several previously unidentified transcription factor targets, approximately 25% of our hits were derived from chromatin-associated proteins, including multiple chromatin remodeling factors, components of lysine modification complexes, and methyllysine readers. These results suggest previously unappreciated mechanisms by which p38 may impact transcription. Furthermore, enrichment of GTPase regulators among JNK1 interactors is interesting in light of the capacity of RHO family GTPases to activate the JNK cascade (*54, 55*), and we verified that the RHO GAPs SYDE1 and SYDE2 are both cellular JNK substrates. JNK kinases may therefore have general roles in crosstalk or feedback regulation between GTPase signaling pathways. Overall, these studies provide a resource for further investigation into regulation of basic cellular processes by the p38 and JNK MAP kinases.

## MATERIALS AND METHODS

### Plasmids

The plasmid for constitutive expression of N-terminally HisMax epitope-tagged human MKK6 in yeast was generated by insertion of the full coding sequence (PCR amplified from pcDNA3-HisMax-MKK6) (*56*) downstream of the *ACT1* promoter (PCR amplified from pGS62, a gift from Gavin Sherlock) in pRS416. MKK6^ΔD^ and D-site substitution mutants on this background were generated by overlap extension PCR using mutagenic oligonucleotides to delete residues spanning Ser4 – Lys17 (SKGKKRNPGLKI) or replace them with D-sites from MKK7 (PQRPRPTLQLPLAN), MEF2A (SRKPDLRVVIPPS) or NFAT4 (LERPSRDHLYLPLE). To generate integrative expression vectors for WT MKK6 and MKK6^D7^, coding sequences were first subcloned into pRS416-GPD downstream of the yeast *TDH3* (GPD) promoter, and then the entire expression cassette was PCR amplified and cloned into the PacI and BglII sites of the plasmid HO-hisG-URA3-hisG-poly-HO (Addgene plasmid #51661). The integrating inducible yeast expression vectors for N-terminally FLAG-tagged human JNK1 (isoform α_1_) and rat p38α were generated by inserting the full-length coding sequences into pRS403-GAL1. The inducible yeast GST expression vector was generated by PCR amplifying the GST coding sequence from pGEX-4T1 and cloning into the SacI and SpeI sites of pRS416-GAL1. Coding sequences for individual D-sites together with the HisMax tag were then PCR amplified from the corresponding pRS416-MKK6 plasmid and inserted into the SpeI and ClaI sites downstream of GST.

Bacterial expression vectors for GST-JNK1 (pGEX4T1-3xFLAG-JNK1, Addgene #47574), GST-p38α, His_6_-MKK4, His_6_-MKK6^S207E/T211E^ (MKK6-EE), and constitutively active MEKK1 (MEKK-C) were previously reported (*57, 58*). Expression vectors for NFAT4 D-site variants were prepared by subcloning residues 3 – 407 of WT NFAT4 from the corresponding mammalian expression vector (Addgene #21664) into pGEX4T1, introducing ClaI and HindIII restriction sites flanking the D-site by site-directed mutagenesis, excising the native D-site coding sequence, and replacing it with synthetic oligonucleotide pairs harboring compatible ends.

The mammalian expression vector for human SYDE1 (Uniprot isoform 2, Q6ZW31-2) was generated by Gateway recombination from pDONR223-SYDE1 (human ORFeome collection) into the C-terminal 3xFLAG epitope tagged plasmid pV1900. The expression vector for N-terminally FLAG-tagged mouse SYDE2 was generated by PCR amplification of the coding sequence from pNICE HA-mSYD1B (Addgene #59362) and insertion into pcDNA3-FLAG by Gibson assembly.

Point mutations in all plasmids were introduced by QuikChange site-directed mutagenesis.

### Design and generation of D-site libraries

The positional scanning libraries consisted of all possible single amino acid substitutions and all double Ala mutations to the MKK6 (SKGKKRNPGLKIPK) and second MKK7 (PQRPRPTLQLPLAN) D-site. To design the proteomic library, we identified all sequences matching the regular expression [RK]-x_0-2_-[RK]-x_3-5_-[ILV]-x-[FILMV] within the human proteome. Sequences were extended at both termini to include two residues downstream of the motif and to bring the total length to 14 residues, and overlapping sequences were removed. Non-cytoplasmic sequences and those falling within annotated PFAM domains were excluded from the final library. Sequences were reverse translated *in silico* to yeast optimized codons, and silent mutations were introduced to remove restriction sites used for cloning. Common flanking sequences were added for separate PCR amplification of the positional scanning (5’: GCTTCAGGTGGACAACAATCACAA, 3’: GAAGCTTCACTCTGTGTTGAAGTTCCGTCAG) and proteomic (5’: GGTCGCGGATCTATGTCTCAG, 3’: GAAGCTTTTGAACAACCTCAGCAC) libraries. Core DNA and protein sequences are provided in Data file S1. Oligonucleotides were commercially synthesized as a pool (CustomArray), PCR-amplified, and restriction enzyme cloned into the NheI and HindIII sites downstream of the GST coding sequence of the pRS416-GAL1-GST plasmid (see Fig. S4 for protein sequence). DH10β cells (Invitrogen ElectroMAX) were transformed by electroporation with ligation products to produce at least 1000 transformants per library variant, and plasmid library DNA was prepared from pooled colonies. To ensure full representation of all components of the library, the variable region was PCR-amplified and sequenced on an Illumina HiSeq 4000 instrument.

### Yeast growth assays

Liquid cultures of the indicated strains transformed with the indicated plasmids were grown to mid-logarithmic phase in the appropriate selective media containing 2% raffinose. Aliquots of five-fold dilution series were spotted onto agar plates containing either 2% glucose or 2% raffinose + 1% galactose as indicated. Plates were incubated at 30℃ for 48 – 96 hours.

### Yeast-based screens

The *S. cerevisiae hog1Δ pbs2Δ* strain was generated by PCR-based replacement of the entire *HOG1* open reading frame with the *LEU2* marker in a *pbs2Δ*::KanMX strain from the yeast knockout collection (BY4741 strain background, Open Biosystems). The genotype was confirmed by diagnostic PCR of both deletion arms from genomic DNA. Strains used for screening were generated by subsequent integration of cassettes for galactose-inducible expression of p38α^L195A^ or JNK1^L198A^ (at the *HIS3* locus) and constitutive GPD promoter-driven expression of His-tagged WT MKK6 or MKK6^D7^ (at the *HO* locus). Expression of MAPK and MKK6 alleles were confirmed by immunoblotting lysates from galactose-treated cells with antibodies to the FLAG and His_6_ tags, respectively.

Libraries of plasmids expressing D-site GST fusion proteins were introduced into yeast by LiOAc high-efficiency transformation (*59*) and selection on SC-Ura agar plates to produce at least 200 transformants per component. Transformed yeast were scraped from plates, pooled, diluted to an OD_600_ of 0.1 in SC-Ura liquid media with 2% glucose, and grown to saturation at 30°C. Cells were diluted into SC-Ura with 2% raffinose and grown for 6 hours to derepress the *GAL1* promoter. A starting time (T_0_) sample (20 OD_600_ units) was reserved, and the remaining cells were split and diluted to an OD_600_ of 0.1 in either SC-Ura + 2% raffinose + 1% galactose (inducing conditions) or SC-Ura + 2% glucose (control conditions). Cultures were subjected to four growth and dilution cycles in which they were propagated until the induced culture reached an OD_600_ of ~1.5, a portion (20 OD_600_ units) reserved, and remaining cells diluted in fresh pre-warmed media to an OD_600_ of 0.1. Reserved cells were pelleted, washed once with sterile dH_2_O, snap-frozen on dry ice/EtOH and stored at −80°C. Plasmids were extracted from each cell pellet and the D-site regions were PCR amplified, incorporating barcodes specific to each condition and time point and adaptors for sequencing. PCR products were agarose gel-purified, pooled, and subjected to Illumina sequencing (HiSeq 4000). The positional scanning library was screened twice, and the human proteomic library was screened three times against each MAPK.

Data were normalized to the total read counts for a given timepoint. Data for each sequence were fit to an exponential function in Microsoft Excel, where the inverse time constant λ was calculated as the slope of the line of the log_2_ transformed fold-change in normalized read counts as a function of time, with the y-intercept set to zero. Z-scores for each sequence within an individual screen were calculated from (λ_sequence_ -λ_mean_)/SD. For the human proteomic screen, p-values were calculated by comparing the Z-scores for a given sequence against the Z-scores for all sequences in the library. The hit threshold (Z ≥ 2, *p* ≤ 0.1) was chosen to maximize the number of true positives while excluding all true negatives.

### Database searches

For PSSM searching, hit sequences were binned into four categories, accounting for sites to occur in a different register from our original definition. JIP class: all JNK1 hits containing a R-P-x-x-ϕ sequence starting at either position 4 or position 6. NFAT4 class: JNK1 hits with an ϕ-x-ϕ-x-ϕ sequence starting at either position 8 or position 10 that were not included in the JIP class. MEF2 class: p38α hit sequences with either a Leu or Pro residue at position 8 or a Pro residue at position 13. Other p38: p38α hit sequences with a Lys or Arg residue at either position 8 or 9, excluding sequences defined in the MEF2 motif. Sequences belonging to each motif class were entered into the program PSSMsearch (http://slim.icr.ac.uk/pssmsearch/) (*32*) and the resulting PSSM used to searched the human proteome with default settings (disorder cutoff = 0.4, p-value cutoff = 0.001). Search results were ranked based on the PWM score, and top 200 sequences are shown in Table S4.

Gene set enrichment analysis for gene ontology categories associated with hit sequences was performed using the Broad Institute web interface (https://www.gsea-msigdb.org/).

### Protein Expression and Purification

GST-tagged (JNK1, p38α, and NFAT4^3-407^ variants) and His_6_-tagged (MKK6-EE and active MKK4 prepared by co-expression with MEKK-C) were expressed in BL21(DE3) *E. coli* and purified as described (*57, 58*). FLAG epitope-tagged SYDE1 and SYDE2 were expressed in and purified from polyethyleneimine-transfected HEK293T cells as previously described (*60*). The concentration and purity of protein preparations were assessed by SDS-PAGE and Coomassie Brilliant Blue R250 staining alongside BSA standards.

GST-p38α and GST-JNK1 (50 μM) were activated *in vitro* by incubation with 500 nM His_6_-MKK6-EE or 50 nM active His_6_-MKK4, respectively, in reaction buffer (50 mM Tris [pH 8.0], 100 mM NaCl, 10 mM MgCl_2_, 1mM DTT, 0.012% Brij-35, 300 μM ATP) at 30°C for 1.5 hours. Phosphorylation was confirmed by immunoblotting with anti-p38 pThr180/pTyr182 and JNK pThr183/pTyr185 antibodies as appropriate.

### Immunoblotting

Samples for immunoblotting were fractionated by SDS-PAGE and transferred to polyvinylidene difluoride membranes. Membranes were blocked with 5% non-fat dry milk in Tris buffered saline with 0.05% Tween-20 (TBS-T) at room temperature for 1 hour and probed with primary antibodies diluted according to the manufacturer’s recommendation. The following primary antibodies used were obtained from Cell Signaling Technology: p38 pThr180/pTyr182 (#9211), c-JUN (#2315), c-JUN pSer63 (#9261), GST (#2624). Other antibodies used were: NFAT4 pSer165 (Sigma-Aldrich SAB4503947), FLAG M2 (Sigma-Aldrich F3165), vinculin (Sigma-Aldrich V9131), penta-His (Qiagen). Membranes were then incubated with fluorophore-conjugated secondary antibodies diluted 1:20,000 in TBS-T and 5% BSA. The fluorescence signal was detected using an Odyssey CLx imaging system (LI-COR Biosciences) and quantified using Image Studio Lite software.

### MAPK D-site peptide inhibition assays

D-site peptides were commercially synthesized (GenScript) incorporating a fixed Tyr-Ala sequence upstream of 14 residues corresponding to the yeast library sequence. Peptides were dissolved in DMSO to 10 mM and stored at −20°C. MAPK assays were performed with a sulfonamido-oxine (SOX) containing substrate peptide (AssayQuant AQT0376). Kinase assays were performed in technical duplicate in black 384 well plates in reactions containing 50 mM HEPES (pH 7.5), 10 mM MgCl_2_, 0.012% Brij-35, 1% glycerol, 0.2 mg/ml BSA, 1 mM ATP, 1.2 mM DTT and 4 μM SOX peptide substrate. Competitor D-site peptides were titrated in two-fold increments over a range from 31 nM – 64 μM. Reactions were initiated by adding activated p38α or JNK1 to final concentrations of 3 nM and 60 nM, respectively, and fluorescence (excitation 360 nm, emission 485 nm) was read every min over 1 hour in a Molecular Devices SpectraMax M5 plate reader. Initial velocities were calculated from the linear portions of the reaction progress curves. IC_50_ values and 95% CIs were calculated by fitting data collected from three biological replicates to a sigmoidal dose-response curve using Prism 8.2.0 (GraphPad).

### Protein kinase assays

GST-NFAT4^3-407^ and its variants (0.5 μM) were incubated with p38α (14 nM) or JNK1 (7 nM) in reaction buffer (50mM HEPES [pH 7.4], 100mM NaCl, 10mM MgCl_2_, 0.012% Brij-35, 1mM DTT, 1mM Na_3_VO_4_, 5mM β-glycerophosphate, 100 μM ATP) at 30°C for 20 min. Reactions were quenched by adding SDS-PAGE loading buffer and analyzed by immunoblotting with antibodies to GST and NFAT4 pSer165.

Purified SYDE1 (100 nM) or SYDE2 (120 nM) was incubated with or without active JNK1 (70 nM) in kinase reaction buffer containing 50 mM HEPES (pH 7.4), 100 mM NaCl, 10 mM MgCl_2_, 0.012% Brij-35, 1 mM DTT, 1 mM Na_3_VO_4_, 1 mM β-glycerophosphate and 50 nM staurosporine (to suppress background phosphorylation in the control reaction). Reactions were initiated by adding [γ-^32^P]ATP to a final concentration of 20 μM at 0.1 μCi/μl. Reactions were incubated at 30°C for 20 min and then quenched with the addition of 5 μl 4x SDS-PAGE loading buffer. Samples were separated by SDS-PAGE (10% acrylamide) and gels were stained with Coomassie, destained, and exposed to a phosphor screen. Exposures were analyzed by phosphorimager and quantified using QuantityOne software (BioRad). Experiments were performed at least three times.

### Analysis of protein phosphorylation in cultured cells

HEK293T cells were transiently transfected with FLAG-tagged SYDE1 and SYDE2 expression plasmids using polyethyleneimine. After 48 h, cells were treated with either 5 μM JNK-IN-8 (SelleckChem, S4901) or vehicle (0.1% DMSO) for 1 hour followed by either 10 μg/mL anisomycin or vehicle (0.1% DMSO) for an additional hour at 37°C. Lysates were prepared as described, and a portion was subjected to either standard or Phos-Tag SDS-PAGE (*61*) followed by immunoblotting with antibodies to the FLAG epitope, total c-JUN, c-JUN pSer63, total JNK, phospho-JNK, and vinculin (all at 1:1000 dilution). Phos-Tag gels included 7.5% acrylamide, 50 μM Phos-Tag reagent (Nard Institute AAL-107), and 100 μM MnCl_2_. incubated in the dark 1 hour with fluorescently-labeled secondary antibodies diluted 1:20,000 in 5% non-fat milk in TBST. Membranes were washed with TBST on rocker at RT 3×10 minutes and imaged with Odyssey CLx (LI-COR Biosciences).

For MS analysis, FLAG-tagged proteins were isolated from 10 cm plates as described above and fractionated by SDS-PAGE. Gels were stained briefly with Coomassie Brilliant Blue and de-stained. Protein bands were excised and subjected to in-gel trypsin digestion and LC-MS/MS analysis at the Yale Keck Biotechnology Resource Laboratory.

## SUPPLEMENTARY MATERIALS

**Figure S1. Growth suppression from co-expression of MKK6 and p38α in yeast.**

**Figure S2. Yeast competitive growth screens.**

**Figure S3. Structures of p38α and JNK1 in complex with D-site peptides.**

**Figure S4. Amino acid sequences of GST fusion libraries.**

**Data file S1. Amino acid and encoding nucleotide sequences for all components of the positional scanning and proteomic libraries.**

**Data file S2. Raw sequencing data for combinatorial library screens.**

**Data file S3. Z-scores for combinatorial library screens.**

**Data file S4. Raw sequencing data for proteomic library screens.**

**Data file S5. Z scores for proteomic library screens.**

**Table S1. List of previously reported functional D-sites present in the proteomic library.**

**Table S2. Hits from proteomic library screens.**

**Table S3. IC_50_ values for D-site peptide inhibition of MAPK activity.**

**Table S4. PSSMsearch results.**

## ACKNOWLEDGMENTS

We thank Elias Lolis, Anton Bennett, Joel Sexton, and Erik Schaefer for advice and suggestions regarding this work, and we thank Titus Boggon for advice and feedback on the manuscript. We thank Karl Barber for assistance with oligonucleotide library design, and TuKiet Lam for LC-MS/MS analysis. We thank Ana Thevenin and Gavin Sherlock for providing plasmids, and the following investigators from whom plasmids were obtained through Addgene: Anjana Rao, Peter Scheiffele, David Stillman and Kevin Janes. This work was supported by National Institutes of Health grant R01 GM135331 to B.E.T. Additional support was provided by the China Scholarship Council to G.S., a National Science Foundation Graduate Research Fellowship to J.T.R. and NIH T32 GM007324 to C.S.

