## Supplemental material for "Proteome-wide screening for mitogen-activated protein kinase docking motifs and interactors"

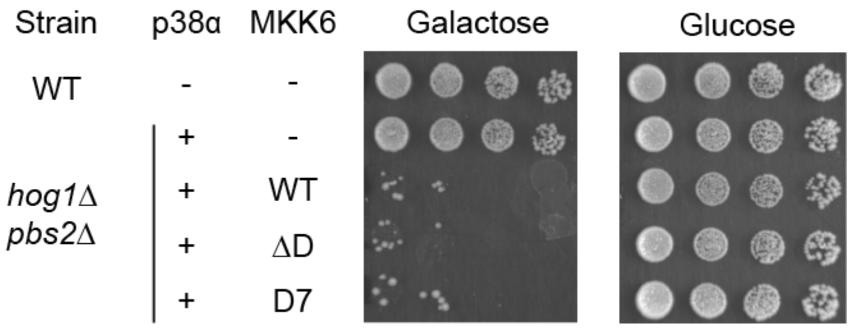


**Figure S1. Growth suppression from co-expression of MKK6 and p38α in yeast.** Assay shows growth inhibition upon co-expression of p38α and the indicated MKK6 variants. Cells of the indicated genotype were spotted in 5-fold serial dilutions on media containing either galactose (to induce p38 expression) or glucose. Growth suppression in response to p38α induction required MKK6 co-expression but was independent of the MKK6 D-site. The capacity of p38α to autophosphorylate in yeast could promote toxic hyperactivation in response to a low MKK6 signal, causing D-site independence.

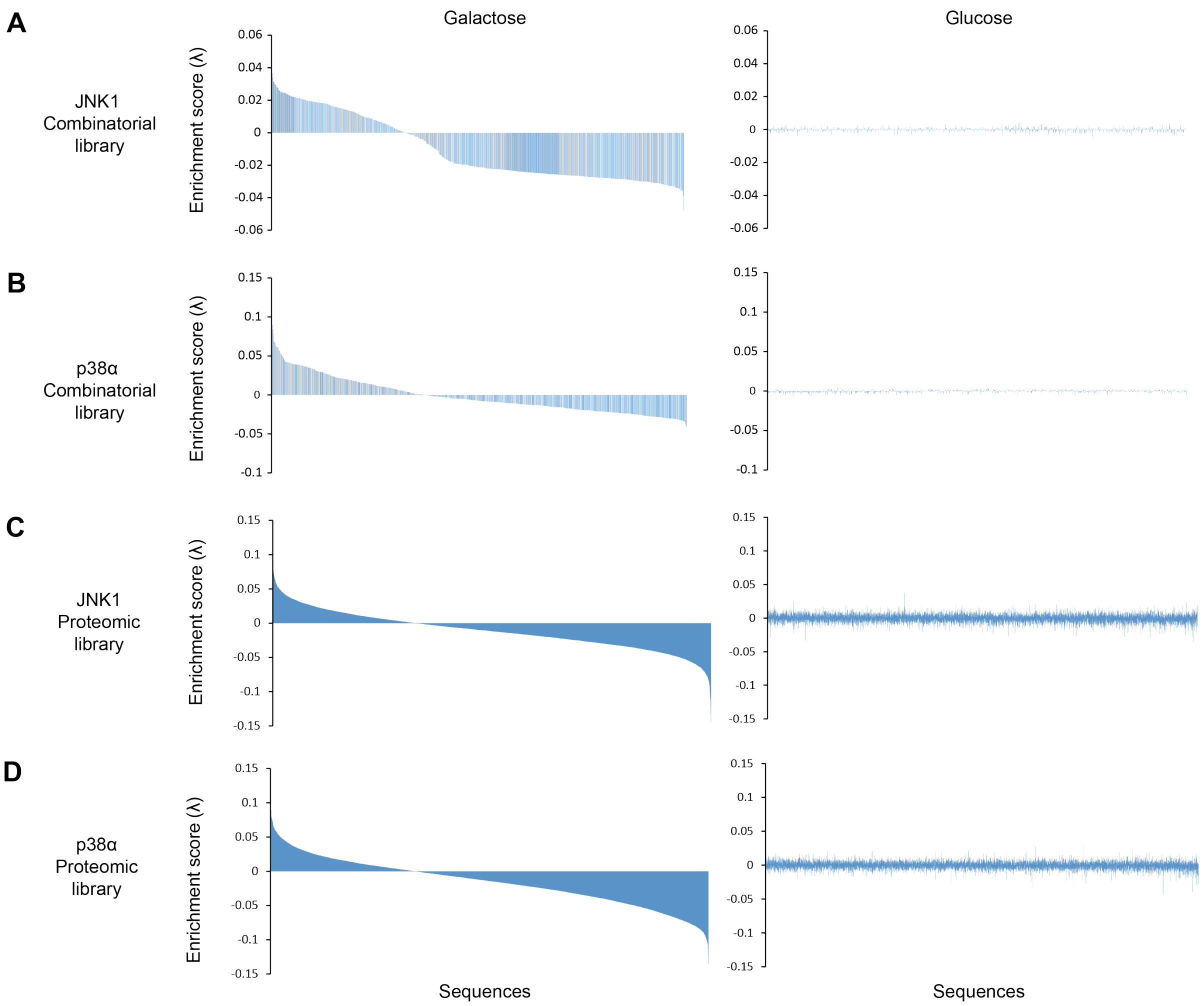


**Figure S2. Yeast competitive growth screens.** Waterfall plots show the enrichment score for each variant in the indicated library upon induction of the indicated MAPK. Sequences are ordered from most enriched to most depleted in galactose within a given screen. Graphs on the right have sequences in the same order but show the enrichment score in the presence of glucose. Representative plots from one of two (combinatorial library) or three (proteomic library) independently performed replicates are shown.

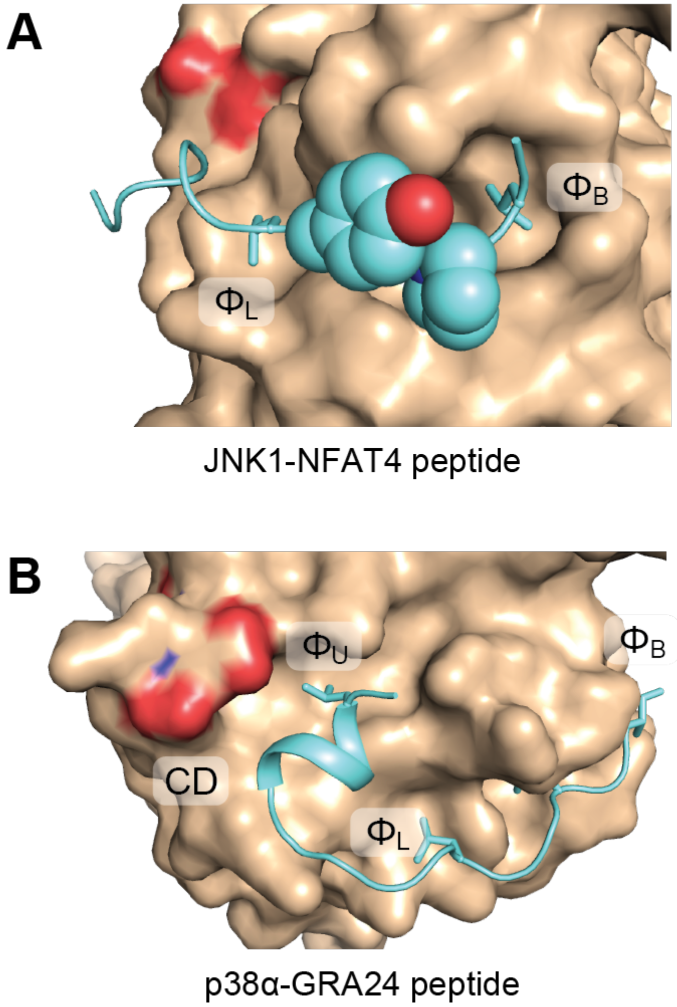


**Figure S3. Structures of p38α and JNK1 in complex with D-site peptides.** (A) Structure of JNK1 in complex with the D-site from NFAT4 (RPSRDHLYLPLE, PDB code 2XS0). Interacting Tyr and Pro residues intervening the $\phi_L$, $\phi_A$, and $\phi_B$ Leu residues are shown as spheres. (B) Structure of p38α in complex with the D-site from *T. gondii* GRA24 (PDB code 5ETA). A Leu residue near the N-terminus of the GRA24 peptide interacts with the hydrophobic $\Phi_U$ pocket proximal to the CD region.

### Positional scanning library GST fusion construct

| | | | | | | | | | | | | | | | | | | |
|---|---|---|---|---|---|---|---|---|---|---|---|---|---|---|---|---|---|---|
| *L* | *V* | *P* | *R* | T | S | T | M | G | S | S | H | H | H | H | H | H | S | S |
| *…CTG* | *GTT* | *CCG* | *CGT* | ACT | AGT | ACC | ATG | GGC | AGC | AGC | CAT | CAT | CAT | CAT | CAT | CAC | AGC | AGC |
| G | L | V | P | R | G | S | H | M | A | S | M | T | G | G | Q | A | S | G |
| GGC | CTG | GTG | CCG | CGC | GGC | AGC | CAT | ATG | GCT | AGC | ATG | ACT | GGT | GGA | CAG | GCT | TCA | GGT |
| G | Q | Q | S | Q | **X** | **X** | **X** | **X** | **X** | **X** | **X** | **X** | **X** | **X** | **X** | **X** | **X** | **X** |
| GGA | CAA | CAA | TCA | CAA | **NNN** | **NNN** | **NNN** | **NNN** | **NNN** | **NNN** | **NNN** | **NNN** | **NNN** | **NNN** | **NNN** | **NNN** | **NNN** | **NNN** |
| E | A | F | E | Q | P | * | | | | | | | | | | | | |
| GAA | GCT | TTT | GAA | CAA | CCT | TAA | | | | | | | | | | | | |

### Proteomic library GST fusion construct

| | | | | | | | | | | | | | | | | | | |
|---|---|---|---|---|---|---|---|---|---|---|---|---|---|---|---|---|---|---|
| *L* | *V* | *P* | *R* | T | S | T | M | G | S | S | H | H | H | H | H | H | S | S |
| *…CTG* | *GTT* | *CCG* | *CGT* | ACT | AGT | ACC | ATG | GGC | AGC | AGC | CAT | CAT | CAT | CAT | CAT | CAC | AGC | AGC |
| G | L | V | P | R | G | S | H | M | A | S | M | T | G | G | Q | Q | G | R |
| GGC | CTG | GTG | CCG | CGC | GGC | AGC | CAT | ATG | GCT | AGC | ATG | ACT | GGT | GGA | CAG | CAA | GGT | CGC |
| G | S | M | S | Q | **X** | **X** | **X** | **X** | **X** | **X** | **X** | **X** | **X** | **X** | **X** | **X** | **X** | **X** |
| GGA | TCT | ATG | TCT | CAG | **NNN** | **NNN** | **NNN** | **NNN** | **NNN** | **NNN** | **NNN** | **NNN** | **NNN** | **NNN** | **NNN** | **NNN** | **NNN** | **NNN** |
| E | A | F | E | Q | P | * | | | | | | | | | | | | |
| GAA | GCT | TTT | GAA | CAA | CCT | TAA | | | | | | | | | | | | |

**Figure S4. Amino acid sequences of GST fusion libraries.** Upstream sequence not depicted is the GST open reading frame from pGEX-4T1, with the C-terminal thrombin cleavage site sequence indicated in italics. The core 14 residue sequence unique to each component of the libraries is indicated in boldface. Residual linker sequence from the synthetic oligonucleotide pool is underlined.

**Table S1. List of previously reported functional D-sites present in the proteomic library.** The starting residue of the two separate sites in BMPR2 are indicated in parenthesis.

| Gene | Protein | Sequence | MAPK | p38 rank | JNK rank |
|---|---|---|---|---|---|
| MAP2K6 | MKK6 | SKGKKRNPGLKIPK | p38 | 17 | |
| MEF2A | MEF2A | MNSRKPDLRVVIPP | p38 | 36 | |
| MAP2K3 | MKK3 | KGKSKRKKDLRISC | p38 | 59 | |
| MEF2C | MEF2C | MNNRKPDLRVLIPP | p38 | 67 | |
| PTPN7 | HePTP | RLQERRGSNVALML | p38 | 238 | |
| PTPN5 | STEP | GLQERRGSNVSLTL | p38 | 434 | |
| MAP2K4 | MKK4 | MQGKRKALKLNFAN | JNK + p38 | 75 | 4 |
| ATF2 | ATF2 | VHKHKHEMTLKFGP | JNK + p38 | | 679 |
| MAPK8IP1 | JIP1 | TYRPKRPTTLNLFP | JNK | | 2 |
| NFATC3 | NFAT4 | ERPSRDHLYLPLEP | JNK | | 3 |
| BMPR2 | BMPR2 (753) | QNLPKRPTSLPLNT | JNK | | 5 |
| MAPK8IP2 | JIP2 | EPHKHRPTTLRLTT | JNK | | 14 |
| BMPR2 | BMPR2 (934) | PRRAQRPNSLDLSA | JNK | | 21 |
| IRS2 | IRS2 | RGRAVRPTRLSLEG | JNK | | 23 |
| MAPK8IP3 | JIP3 | GRRKERPTSLNVFP | JNK | | 26 |
| DUSP10 | MKP5 | LSRPVRPQDLNLCL | JNK | | 30 |
| IRS1 | IRS1 | NSRLARPTRLSLGD | JNK | | 37 |
| MAP2K7 | MKK7 | PQRPRPTLQLPLAN | JNK | | 38 |
| SPAG9 | JIP4 | RIRKERPISLGIFP | JNK | | 85 |

**Table S3. $IC_{50}$ values for D-site peptide inhibition of MAPK activity.**

| | | JNK1 | | p38α | |
|---|---|---|---|---|---|
| Gene Name | Peptide sequence | $IC_{50}$ (μM) | 95% CI range | $IC_{50}$ (μM) | 95% CI range |
| ARHGEF5 | LRRKLNTRPVHLHL | 2.4 | 2.0 - 2.8 | 16 | 11 - 24 |
| ATP8B1 | KTKRNKILKLKFPR | 6.8 | 5.4 - 8.4 | 1.4 | 1.1 - 1.8 |
| CCAR2 | LRRRLTPLQLEIQR | 3.9 | 3.2 - 4.8 | 24 | 21 - 28 |
| CCNB3 | EGKRSRLKPLVLQE | 4.5 | 3.8 - 5.3 | 92 | 79 - 108 |
| CMYA5 | KKGVKPKLVLNVTS | 12 | 9.5 - 15.4 | 14 | 11 - 19 |
| DOCK6 | ATVKARVAELYLPL | 4.1 | 3.4 - 4.9 | 24 | 17 - 34 |
| DOCK8 | PEVKVKIAALYLPL | 5.7 | 4.5 - 7.2 | 14 | 10 - 20 |
| EIF2B5 | KSKWCRPTSLNVVR | 0.40 | 0.36 - 0.45 | >100 | - |
| ELMSAN1 | PKQRPRPEPLIIPT | 58 | 47 - 73 | 74 | 54 - 105 |
| FAM214B | GRRLKGARRLKLSP | 45 | 39 - 53 | 6.0 | 4.8 - 7.6 |
| KIF20B | INEKKEKLTLEFKI | 13 | 9.9 - 18 | >100 | - |
| L3MBTL3 | KCSRKKKPKLSLKA | 29 | 23 - 36 | 3.0 | 2.239 - 4.145 |
| LRCH4 | GEERRRPDTLQLWQ | 0.76 | 0.66 - 0.88 | >100 | - |
| MGA | PGKRGRPRKLKLCK | 1.2 | 0.96 - 1.6 | 2.6 | 2.2 - 3.2 |
| MLXIPL | SPKWKNFKGLKLLC | 12 | 9.8 - 13.5 | 1.0 | 0.74 - 1.5 |
| NCOR2 | NQAMRKKLILYFKR | 0.32 | 0.28 - 0.35 | 4.3 | 4.0 - 4.8 |
| OBSL1 | HRHRLVLNGLGLAD | 10.2 | 8.3 - 12.5 | >100 | - |
| PRAME | EKVKRKKNVLRLCC | 30 | 25.9 - 34.6 | 7.7 | 6.2 - 9.6 |
| PSME4 | KQLKRTHKKLTINP | >100 | - | 42 | 31 - 58 |
| SETD1B | NQLKFRKKKLKFCK | 48 | 38 - 63 | 4.5 | 3.4 - 6.0 |
| SMARCD1 | PIKQKRKLRIFISN | 22 | 18 - 27 | 2.9 | 2.3 - 3.6 |
| SYDE2 | RQKKERPHMLNLSG | 1.0 | 0.93 - 1.1 | >100 | - |
| TRERF1 | KKFRHRPEPLFIPP | 52 | 46 - 60 | 5.2 | 4.5 - 5.9 |
| KMT2D | RKTDRPALHLRIPP | 9.5 | 8.6 - 10.5 | 2.3 | 2.1 - 2.5 |
| KMT2D D4R | RKTRRPALHLRIPP | 2.5 | 1.9 - 3.4 | 0.2 | 0.20 - 0.27 |
| KMT2D I12F | RKTDRPALHLRFPP | 8.0 | 6.9 - 9.2 | 15 | 14 - 17 |
| KMT2D R11E | RKTDRPALHLEIPP | 14 | 11 - 17 | 5.8 | 4.8 - 6.9 |
| KMT2D I12L/P13A | RKTDRPALHLRLAP | 4.6 | 3.077 - 6.89 | 6.4 | 5.2 - 8.0 |

**Additional files**

**Table S2. Hits from proteomic library screens.**

**Table S4. PSSMsearch results.**

**Data file S1. Amino acid and encoding nucleotide sequences for all components of the positional scanning and proteomic libraries.**

**Data file S2. Raw sequencing data for combinatorial library screens.**

**Data file S3. Z-scores for combinatorial library screens.**

**Data file S4. Raw sequencing data for proteomic library screens.**

**Data file S5. Z scores for proteomic library screens.**

**Data file S6. MS data for SYDE2 isolated from HEK293T cells.**
